# Oscillatory signatures of crossmodal congruence effects: An EEG investigation employing a visuotactile pattern matching paradigm

**DOI:** 10.1101/014092

**Authors:** Florian Göschl, Uwe Friese, Jonathan Daume, Peter König, Andreas K. Engel

**Author notes:** **Correspondence address:** Florian Göschl, Department of Neurophysiology and Pathophysiology, University Medical Center Hamburg-Eppendorf, Martinistr. 52, 20246 Hamburg, Germany.

## Abstract

Coherent percepts emerge from the accurate combination of inputs from the different sensory systems. There is an ongoing debate about the neurophysiological mechanisms of crossmodal interactions in the brain, and it has been proposed that transient synchronization of neurons might be of central importance. Oscillatory activity in lower frequency ranges (< 30 Hz) has been implicated in mediating long-range communication as typically studied in multisensory research. In the current study, we recorded high-density electroencephalograms while human participants were engaged in a visuotactile pattern matching paradigm and analyzed oscillatory power in the theta- (4-7 Hz), alpha-(8-13 Hz) and beta-bands (13-30 Hz). Employing the same physical stimuli, separate tasks of the experiment either required the detection of predefined targets in visual and tactile modalities or the explicit evaluation of crossmodal stimulus congruence. Analysis of the behavioral data showed benefits for congruent visuotactile stimulus combinations. Differences in oscillatory dynamics related to crossmodal congruence within the two tasks were observed in the beta-band for crossmodal target detection, as well as in the theta-band for congruence evaluation. Contrasting ongoing activity preceding visuotactile stimulation between the two tasks revealed differences in the alpha- and beta-bands. Source reconstruction of between-task differences showed prominent involvement of premotor cortex, supplementary motor area, somatosensory association cortex and the supramarginal gyrus. These areas not only exhibited more involvement in the pre-stimulus interval for target detection compared to congruence evaluation, but were also crucially involved in post-stimulus differences related to crossmodal stimulus congruence within the detection task. These results add to the increasing evidence that low frequency oscillations are functionally relevant for integration in distributed brain networks, as demonstrated for crossmodal interactions in visuotactile pattern matching in the current study.

## 1. Introduction

Multimodal processing and the integration of information from different sensory systems are crucial for adaptive behavior. Crossmodal interactions have been shown to influence perception (sometimes also resulting in illusory percepts, McGurk and McDonald, 1976) as well as a broad range of cognitive processes (Doehrmann and Naumer, 2005; Driver and Spence, 2000; Macaluso and Driver, 2005; Stein, 2012). Behavioral benefits resulting from the successful combination of inputs from different modalities are well documented, including improvements in target detection, discrimination or localization performance and faster response latencies (Diederich and Colonius, 2007; Forster et al., 2002; Frassinetti et al., 2002; Gillmeister and Eimer, 2007; McDonald et al., 2000; Molholm et al., 2004; Murray et al., 2001).

The question of how interactions between different sensory and other regions of the brain are implemented neurophysiologically remains a matter of dispute. Yet, it is evident that in order to quickly adapt to the environment fast changes in functional brain networks are essential (Jensen and Mazaheri, 2010). One mechanism that has been proposed to address the challenge of integration in distributed networks is transient synchronization of neurons (Singer and Gray, 1995; Engel et al., 2001; Varela et al., 2001; Womelsdorf et al., 2007). Only recently, synchronized oscillatory activity has also been linked to the integration of object features across sensory modalities (e.g., Lakatos et al., 2007; Kayser et al., 2008; for a review see Senkowski et al., 2008). Experimental support for a major role of neuronal oscillations in multisensory integration is available for activity below 30 Hz in the theta-, alpha- and beta-bands (Doesburg et al., 2009; Gleiss and Kayser, 2014a; Hummel and Gerloff, 2005; Senkowski et al., 2006) as well as for high frequency activity above 30 Hz (the gamma-band, see for example Bauer et al., 2009; Doesburg et al., 2008; Schneider et al., 2008; Schneider et al., 2011). Explicitly advocating different roles of high and low frequency activities in the integration of distributed information, van Ackeren and colleagues (2014) have demonstrated for lexical-semantic stimulus material that linking modality-specific feature words to a target word is associated with enhanced gamma-band activity between 80 and 120 Hz, whereas integration of features from different modalities is reflected in low-frequency power increases between 2 and 8 Hz. The authors position their findings within a framework proposed by Donner and Siegel (2011), arguing that low-frequency oscillatory activity is involved in the coordination of distributed neuronal populations, while local encoding happens within higher frequency ranges. This matches results by von Stein et al. (2000) reporting synchronization in the gamma-band for directly connected cortical areas and lower frequency synchronization for large-scale networks.

In order to study the multisensory interplay between vision and touch – a well-suited model for long-range communication in the brain – we employed a matching paradigm requiring the identification of concurrently presented visual and tactile dot patterns. In a behavioral study examining the interdependency of crossmodal stimulus congruence and attention (Göschl et al., 2014), we found that congruent as compared to incongruent visuotactile stimulation reliably led to enhanced behavioral performance, mirrored in higher accuracies and shortened reaction times. In the present study, we used a similar paradigm and additionally recorded high-density electroencephalograms (EEG) to investigate the neurophysiological correlates of these congruence effects. Following the idea described earlier that integrative functions involving long-range interactions are predominantly mediated by lower frequencies (Donner and Siegel, 2011; von Stein et al. 2000; see also Hipp et al., 2011) we focused on oscillatory activity below 30 Hz in our analysis.

The visuotactile matching paradigm used here involved two different tasks. In different blocks of the experiment, participants were either asked to (1) detect predefined target patterns that could appear in both sensory modalities (*detection task*) or, (2) explicitly evaluate the relationship between the two patterns and report whether they were the same or not (*congruence evaluation task*). Keeping the physical stimulation identical in both tasks, we investigated between-task differences associated with distinct cognitive demands of *detection* and *congruence evaluation* in anticipation of crossmodal stimulation, as well as within-task differences related to visuotactile stimulus congruence in the whole post-stimulus interval. Whereas stimulus congruence is only of implicit importance for the *target detection*, it is explicitly relevant for the *evaluation task*. We sought to assess the influence of crossmodal congruence that is either passively perceived (*detection*) or actively searched for (*congruence evaluation*) and expected this distinction to be reflected in oscillatory signatures.

Evidence linking brain oscillations to visuotactile interactions is sparse, which is why we decided to pursue an exploratory data analysis approach in the current study. Accordingly, our a-priori hypotheses were formulated cautiously. On the one hand, we expected modulations in preparatory neuronal activity related to task requirements, especially in alpha- and beta-frequencies (Mazaheri et al., 2014; van Ede et al., 2011; van Ede et al., 2014). On the other hand and in line with previous studies on crossmodal interactions and low frequency modulations (Barutchu et al., 2013; Doesburg et al., 2009; Gleiss and Kayser, 2014a; Gleiss and Kayser, 2014b; Hummel and Gerloff, 2005; van Ackeren et al., 2014; van Driel et al., 2014), we expected post-stimulus modulations below 30 Hz related to crossmodal stimulus congruence. In order to investigate the integration of information in visuotactile networks on the level of cortical sources, we performed EEG source reconstruction using eLORETA.

## 2. Methods

### 2.1 Participants

Sixteen right-handed volunteers (12 female, mean age 25.4, range 21-33) received monetary compensation for their participation in the study. All participants had normal or corrected to normal vision, and reported no history of neurological or psychiatric illness. The Ethics Committee of the Medical Association Hamburg approved the current study, and participants provided written informed consent prior to the recordings.

### 2.2 Task design

The experimental setup outlined in the following is similar to a previous behavioral study (Göschl et al., 2014), using only a subset of the visuotactile matching paradigm described in detail in Göschl et al. (2014) (see Figure 1 for an overview of events and timing of the current experiment). Four spatial patterns, each of them formed by three dots, constituted the stimulus set (Figure 1A). Stimuli were presented visually on a computer monitor, appearing left of a central fixation cross, embedded in a noisy background. Concurrently, dot patterns were delivered haptically to the participants’ right index fingertip via a Braille stimulator (QuaeroSys Medical Devices, Schotten, Germany). Stimulus duration was 300 ms for both patterns.

**Figure 1.**
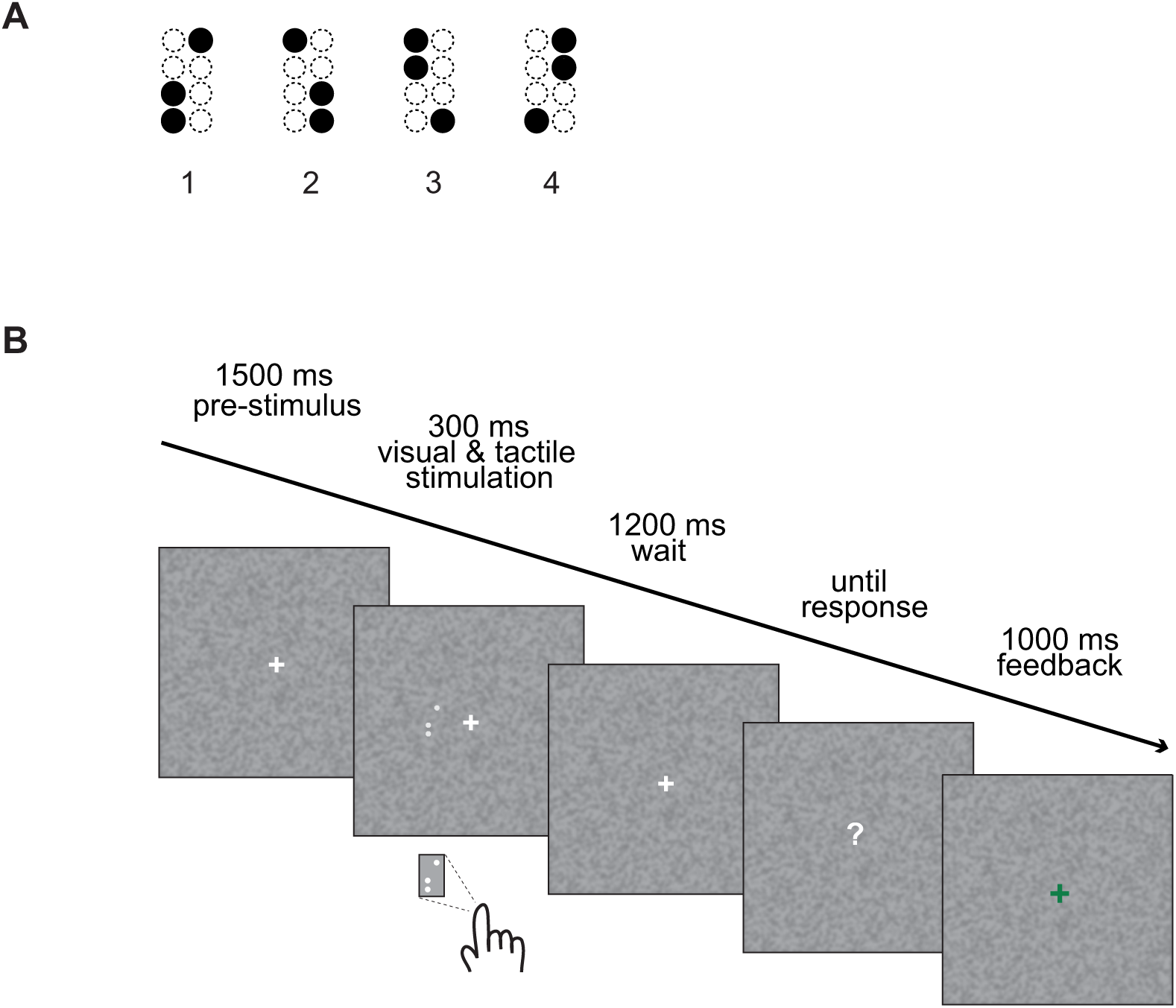
Schematic representation of the visuotactile matching task. **(A)** The four pattern stimuli used in our experiment. **(B)** The trial sequence. After a pre-stimulus interval of 1500 ms, visual and tactile stimuli were presented simultaneously for 300 ms, followed by a wait interval of 1200 ms. After that, a question mark appeared on the screen indicating that responses could be given. After button press, every trial ended with visual feedback (1000 ms).

Prior to the actual experiment, we conducted a delayed-match-to-sample training task to familiarize participants with the tactile patterns. In this training task, participants were asked to judge whether a sample stimulus (duration 300 ms) and a probe stimulus (also of 300 ms duration) presented 1000 ms later were identical or not. Responses were given with the left hand via button press on a response box (Cedrus, RB-420 Model, San Pedro, USA) and visual feedback (a green ‘+’ or a red ‘−’) informed participants about the correctness of their response. After a minimum of five training blocks (each consisting of 16 trials) and a matching performance of at least 80%, participants could proceed to the actual experiment. One participant not meeting this criterion was excluded after the training procedure.

The experimental session incorporated two different tasks (performed in separate blocks of the experiment), which both required the identification of concurrently presented visual and tactile patterns. In the *detection task*, the participants were instructed to detect target stimuli that could appear in both modalities. In each experimental block, one out of the four dot patterns was defined as target (the other three patterns were non-targets, respectively) and introduced to the participants at the start of the block by simultaneously presenting it on the computer screen and by means of the Braille stimulator (four times). In the following experimental trials, targets could appear in the visual or the tactile modality alone, in both or in neither of the two. Participants had to decide whether the presented patterns matched the previously defined target stimulus or not and press one of two response buttons accordingly. In the *congruence evaluation task*, participants were asked to compare patterns across sensory modalities and report whether they were the same (congruent) or not. Again, responses were given via button press.

The timing was identical for the *detection* and the *congruence evaluation tasks* and is displayed in Figure 1B. The major difference compared to the experimental design realized in our earlier study is the wait interval of 1200 ms between stimulus presentation and response. This interval was chosen to prevent contamination of the EEG signal by activity resulting from response execution.

Each participant performed 1536 trials over two sessions recorded on separate days (with the two sessions happening within three days). The experimental design was counterbalanced in the presentation of congruent and incongruent stimulus pairs, target definitions and presentation frequencies of each of the four patterns across trials (for details see Göschl et al., 2014). We pooled data from the two recording sessions and grouped trials as follows: visual targets alone (a visual target appearing with a tactile non-target; labeled incongruent V in the following), tactile targets alone (a tactile target presented with a visual non-target; incongruent T), and visuotactile targets (congruent VT) as well as non-target congruent stimulus pairs and non-target incongruent pairs for the *detection task* (five conditions); for the *congruence evaluation task* we split trials in congruent and incongruent visuotactile stimulus pairs, respectively. This procedure left us with a total of 192 trials for each condition (only the non-target incongruent pairs appeared 384 times to balance tactile and visual target trials). In the following, we focus on correctly detected (incongruent V, incongruent T and congruent VT) targets for the *detection task* and accurately identified congruent and incongruent stimulus pairs for the *congruence evaluation task*.

The mapping of response keys (for ‘target’ and ‘non-target’-, as well as ‘congruent’ and ‘incongruent’-buttons) was counterbalanced across participants and sessions. To mask sounds associated with pin movement in the Braille cells, the participants were presented with pink noise administered via foam-protected air tube earphones at a 75 dB sound pressure level (Eartone, EAR Auditory Systems, AearoCompany). We used Presentation software (Neurobehavioral Systems, version 16.3) to control stimulus presentation and to record participants’ response times (RT) and accuracies.

### 2.3 EEG recordings

EEG data were acquired from 126 scalp sites using Ag/AgCl ring electrodes mounted into an elastic cap (EASYCAP, Herrsching, Germany). Additionally, two electrodes were placed below the eyes to record the electrooculogram. EEG data were recorded with a passband of 0.016-250 Hz and digitized with a sampling rate of 1000 Hz using BrainAmp amplifiers (BrainProducts, Munich, Germany). The tip of the nose served as a reference during the recordings but subsequently we re-referenced the data to common average. Analysis of the EEG data was carried out in Matlab 8.0 (MathWorks, Natick, MA) using custom-made scripts, as well as routines incorporated in EEGLAB 11.0 (Delorme and Makeig, 2004; http://sccn.ucsd.edu/eeglab/) and FieldTrip (Oostenveld et al., 2011; http://fieldtrip.fcdonders.nl). Offline, the data were band-pass filtered (0.3-180 Hz), downsampled to 500 Hz and epoched from − 400 to + 1400 ms around the onset of the simultaneously presented visual and tactile stimuli. All trials were inspected visually, and those containing EMG artifacts were rejected. Afterwards we applied an independent component analysis (ICA) approach to remove artifacts related to eye blinks, horizontal eye movements and electrocardiographic activity. To control for miniature saccadic artifacts, we employed the COSTRAP algorithm (correction of saccade-related transient potentials; Hassler et al., 2011) that has been used to suppress ocular sources of high frequency signals (e.g., Friese et al., 2013; Hassler et al., 2013). With this multilevel artifact correction procedure, 88% of all recorded trials (range: 75% to 95%) were retained.

#### 2.3.1 Spectral analysis

We derived time-frequency representations of the data via wavelet convolution in the frequency domain. Fast Fourier transforms of the EEG signal were obtained and multiplied by the Fourier transform of the complex Morlet wavelets [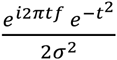, where *t* represents time, *f* is frequency which increased in 30 logarithmic steps from 2 to 100 Hz, and *σ* defines the width of each frequency band, set according to *n/(2πf)*, where *n* stands for the number of wavelet cycles which increased from 3 to 10 in logarithmic steps (Cohen and Donner, 2013; Cohen, 2014)]. Then, the inverse fast Fourier transform was taken. All frequency transformations were done at the single-trial level before averaging. Power estimates for specific frequencies at each time point were defined as the squared magnitude of the complex convolution result {real[z(t)]^2^ + imaginary[z(t)^2^]}. To compute the relative signal change, power data were normalized with respect to a pre-stimulus baseline window. The baseline power was calculated as the average from – 300 ms pre-stimulus to 0 (stimulus onset).

To obtain the induced, non-phase-locked part of the signal power, we computed the event-related potential and subtracted it from the time domain signal on each trial (Kalcher and Pfurtscheller, 1995). This procedure was carried out for each condition, electrode and subject separately. Afterwards, the time-frequency decomposition was conducted as described in the previous paragraph. Analysis was done for both, total and induced power with results being highly comparable. For reasons of clarity, we focus on the analysis of induced power in the following.

Figure 2A and 2B show baseline corrected time-frequency representations averaged across all sensors, experimental conditions (also including the non-target conditions for the *detection task*) and participants for the two tasks. To investigate within-task differences related to crossmodal stimulus congruence as well as between-task differences related to different cognitive demands of *detection* and *congruence evaluation*, we binned and averaged power data in segments of 100 ms covering the whole trial period from −300 to 1300 ms in three frequency bands of interest: theta (4-7 Hz), alpha (8-13 Hz), and beta (13-30 Hz). Poststimulus comparisons within the two tasks were calculated from stimulus onset to 1300 ms after stimulation in steps of 100 ms on baseline-normalized power in all three frequency bands. Figure 2C and 2D show topographies of baseline corrected power for all correct trials of the two tasks averaged for the three frequency bands of interest and time windows that significantly differed from baseline activity (areas showing intense color scaling in Figure 2A and 2B).

**Figure 2.**
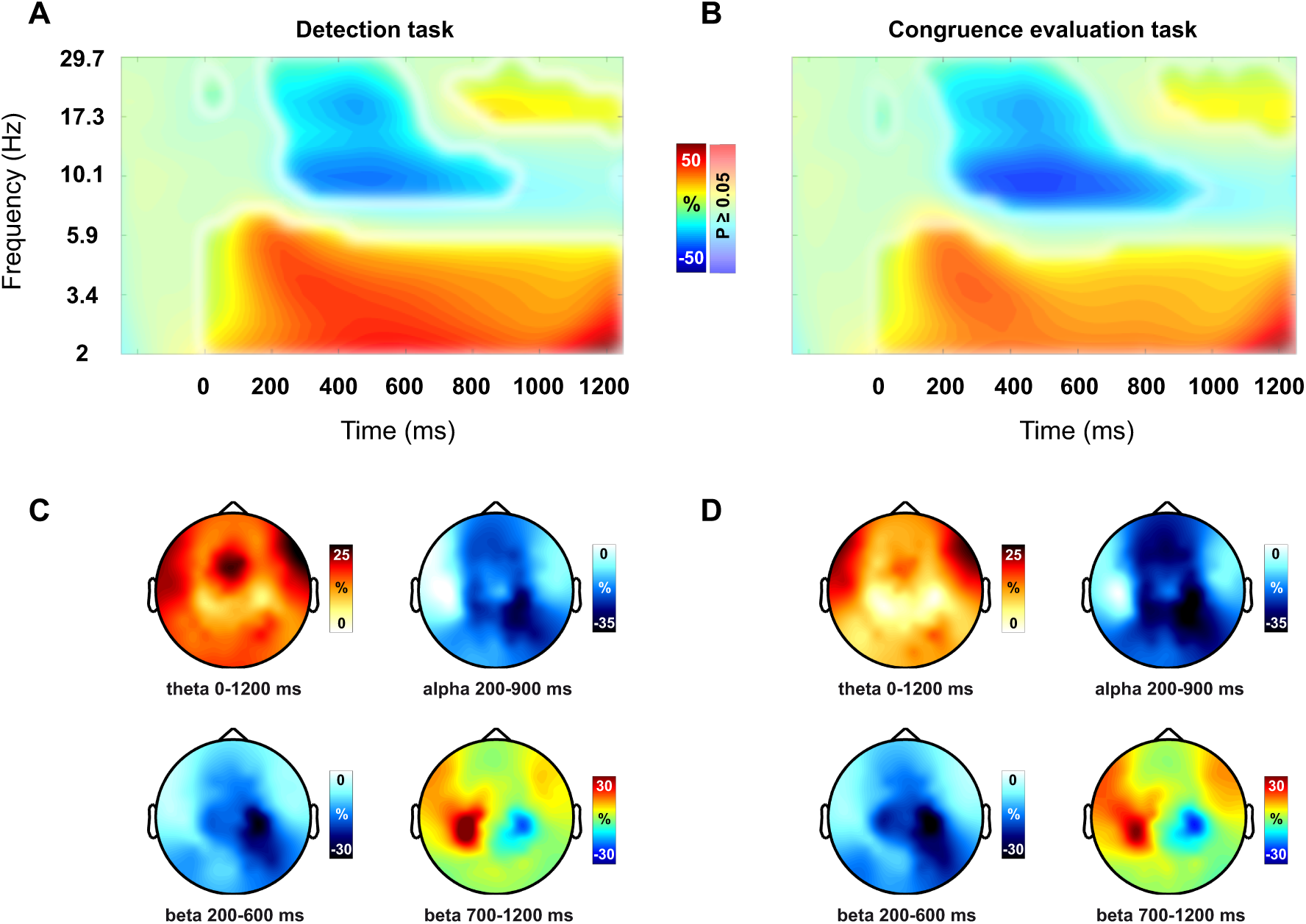
Grand average time-frequency representations for the two tasks and topographies for theta-, alpha- and beta-band power in post-stimulus time windows significantly different from baseline. (**A**) Data for the *detection task*, and (**B**) the *congruence evaluation task*. Power data were averaged across all sensors, experimental conditions (correct trials) and participants and are shown as % change to baseline. Unmasked regions (intense color scale) indicate significant difference to baseline (t-test, p < 0.05, FDR corrected). (**C**) Topographies for power data (% change) significantly different from baseline activity averaged over all correct trials of the *detection task* for theta-band activity between 0 and 1200 ms, alpha-band activity between 200 and 900 ms, beta-band activity between 200 and 600 ms, and beta-band activity between 700 and 1200 ms. (**D**) Power topographies for all correct trials from the *congruence evaluation task* are shown for the same time-frequency windows as in (C).

To assess global differences in oscillatory dynamics between the two tasks, we compared neuronal activity directly preceding the presentation of the visual and tactile stimuli (in steps of 100 ms from – 300 ms to stimulus onset), again in theta-, alpha- and beta-frequencies. Raw power values (no baseline normalization applied) were chosen for pre-stimulus comparisons between the tasks.

For the statistical analysis of sensor level power data, we applied a cluster level randomization approach (Maris and Oostenveld, 2007) as implemented in FieldTrip (Oostenveld et al., 2011). This procedure has been used previously (see for example Jokisch and Jensen, 2007; Nieuwenhuis et al., 2008; Mazaheri et al., 2014) and controls for Type I errors involving multiple comparisons (in our case over multiple time points and sensors). First, data for the different conditions respectively tasks were averaged for the frequency bands of interest and within every time bin and a t-statistic was computed for every sensor. Then, contiguous sensors falling below a p-value of 0.05 were grouped in clusters, with the sum of t-values in a given cluster being used in the cluster-level test statistics. Subsequently, the Monte Carlo estimate of the permutation p-value of the cluster was obtained by evaluating the cluster-level test statistic under the randomization null distribution assuming no condition difference. This distribution was created by randomly reassigning the data to the conditions across participants 1000 times and computing the maximum cluster-level test statistic. Analysis was carried out separately for the three frequency bands of interest. To account for comparisons in multiple frequency bands, we applied Bonferroni correction to the significance level used and report cluster test statistics significant if the corresponding p-value falls below the corrected alpha of 0.0167 (0.05/3). To avoid differences in EEG activity resulting from differences in trial count or signal-to-noise ratio, conditions respectively tasks were trial-matched within every participant before contrasting power.

#### 2.3.2 Source estimation of frequency-specific activity

Neuronal sources of frequency band-specific activity were reconstructed using eLORETA (exact low-resolution brain electromagnetic tomography). eLORETA is a non-adaptive linear spatial filter with the property that single dipoles without additional noise can be localized exactly (for technical details see for example Pascual-Marqui, 2007). We calculated cross-spectral density matrices between all 126 scalp EEG channels (excluding the EOG channels) in every frequency (2 to 100 Hz in 30 logarithmic steps) for sixteen time windows of equal length (100 ms, from – 300 to 1300 ms). Calculation of cross spectra was done separately for every participant and experimental condition in each trial. To derive the source estimates, we multiplied the real part of the frequency domain data (the real part of the cross spectrum) with the real-valued filter. We took the largest eigenvalue of the reduced 3 x 3 cross spectrum as a power estimate for each grid point. eLORETA computations were made in a realistic 3-shell head model based on the MNI152 template brain (Montreal Neurological Institute; http://www.mni.mcgill.ca). Source activity was estimated within a continuous grid of 3000 voxels and leadfields were calculated as described in Nolte and Dassios (2005). Source data were baseline corrected as well, using the interval from – 300 ms to 0 (corresponding to the first three time windows of our cross-spectral density matrix calculation).

Across participants, paired t-tests on source level power data were calculated for time and frequency windows that turned out significant in the sensor level analysis. FDR correction (alpha = 0.1) was applied to correct for comparisons involving multiple locations (voxels) and t-maps were masked accordingly. Depending on the results of the FDR correction, t-values are either displayed with FDR correction masks or uncorrected using a t-threshold of ± 2.1314 (corresponding to the 2.5^th^ and 97.5^th^ percentiles of the t distribution with 15 degrees of freedom). Anatomical labeling was done using the NFRI functions (Singh, Okamoto et al., 2005; http://www.jichi.ac.jp/brainlab/tools.html).

## 3. Results

### 3.1 Behavioral data

To determine whether behavioral performance within the *detection task* differed depending on visuotactile stimulus congruence, we subjected accuracy and reaction time data for congruent VT, incongruent V and incongruent T targets to 1 x 3 repeated measures ANOVAs with Congruence as the within-subject factor. We found a significant effect of Congruence (*F*_2, 30_ = 23.28, *p* < 0.01), with congruent VT targets being associated with the highest detection rate, followed by incongruent V targets and incongruent T targets (see Table 1 for mean accuracies of the different conditions). To further elucidate stimulus-congruence related effects, post hoc t-tests were conducted showing that congruent stimulation (VT targets) led to superior detection as compared to incongruent V targets (*t*_15_ = 4.37, *p* < 0.01) and incongruent T targets alone (*t*_15_ = 6.58, *p* < 0.01; paired sample t-tests). Mean reaction times (note that responses could only be given after a forced wait interval of 1200 ms after stimulus presentation) for the detection of congruent VT, incongruent V and incongruent T targets are also displayed in Table 1. Again, a repeated measures ANOVA revealed a significant effect of Congruence (*F*_2, 30_ = 6.59, *p* < 0.01). Post hoc comparisons (paired sample t-tests) showed that reactions were fastest for the detection of congruent VT targets with a significant difference from incongruent T targets (*t*_15_ = 3. 11, *p* < 0.01) and a trend for faster reactions to congruent VT targets compared to incongruent V targets (*t*_15_ = 1.91, *p* = 0.08). Thus, we did not observe a speed accuracy tradeoff; instead bimodal stimulation achieved consistently higher performance.

**Table 1.** Behavioral data of the visuotactile matching paradigm. Mean accuracies (ACC, shown in percent) and response times (RT, shown in ms), with standard deviations (SD) for the *detection* and the *congruence evaluation task* as well as the comparison of the two tasks.

|  | ACC (SD) | RT (SD) |
| --- | --- | --- |
| *Detection task* |  |  |
| Congruent VT targets | 96.2 (3.6) | 376 (193) |
| Incongruent V targets | 88.5 (9.2) | 390 (194) |
| Incongruent T targets | 71 (16.3) | 411 (193) |
| *Congruence evaluation task* |  |  |
| Matching stimulus pairs | 87.5 (7.6) | 406 (200) |
| Non-matching stimulus pairs | 80.4 (16.3) | 417 (195) |
| *Comparison between detection (also including non-targets) and congruence evaluation task* |  |  |
| Detection | 82.9 (7.3) | 410 (203) |
| Congruence evaluation | 83.9 (10.8) | 412 (197) |

In order to compare behavioral performance on matching and non-matching visuotactile pairs of stimuli in the *congruence evaluation task*, we employed paired sample t-tests. Across subjects, congruent pattern combinations showed a trend for higher accuracy (*t*_15_ = 2.07, *p* = 0.06; see Table 1 for mean accuracies and reaction times). This is compatible with a response bias ‘in doubt towards congruence’. The analogous comparison for reaction times yielded no significant result.

Global differences in behavioral performance between the *detection* and the *congruence evaluation tasks* were assessed by calculating mean accuracies and reaction times within the two tasks (including non-target conditions for the *detection task*) and contrasting values between them (paired sample t-tests, two-tailed). Behavioral metrics were comparable, i.e., no significant differences were found.

### 3.2 EEG data

The following section on the results of our EEG sensor and source level analysis is subdivided into three parts: (1) stimulus-congruence related effects in the *detection task*, (2) differences between matching and non-matching stimulus pairs in the *congruence evaluation task* and, (3) a baseline comparison of the two tasks.

#### 3.2.1 Detection task

To analyze frequency-specific differences related to crossmodal stimulus congruence in the *detection task*, we compared responses to congruent VT and incongruent V targets as well as congruent VT and incongruent T targets separately across the whole post-stimulus trial period (0 to 1300 ms, in steps of 100 ms). Additionally, we reconstructed sources of neuronal activity for the different conditions and calculated source power contrasts within time and frequency ranges that showed statistical difference on the sensor level.

Theta-band power was enhanced throughout the whole trial (see also Figure 2C) as compared to baseline, without showing differences related to visuotactile congruence. Alpha- and beta-band decreases starting at around 200 ms after stimulus onset were also comparable between congruent VT and incongruent V or incongruent T trials, respectively.

Differences in oscillatory dynamics within the detection task were observed in the beta-band starting at around 700 ms after the onset of the two patterns (see Figure 3). Cluster statistics revealed differences for the comparison of congruent VT and incongruent T targets in five time bins from 700 to 1200 ms (*p* = 0.006). Figure 3A shows topographies of the evolvement of these stimulus congruence related effects over time starting in left-hemispheric central regions and spreading towards the midline and right-hemispheric central regions (asterisks mark the channels belonging to the cluster showing significant statistical difference). Source level maxima for the comparison of congruent VT and incongruent T target trials (with power values averaged for the interval from 700 to 1200 ms, see Figure 3B) were located in right-hemispheric premotor cortex, supplementary motor area (SMA) and supramarginal gyrus. Similarly, the comparison of beta-band responses to congruent VT and incongruent V trials yielded significant effects in a time window from 700 to 1200 ms. A significant cluster (*p* = 0.007, Figure 3C) is apparent in right-hemispheric posterior and central scalp regions. Maxima of source level differences for the whole time interval were found in right-hemispheric somatosensory association cortex and supramarginal gyrus (Figure 3D).

**Figure 3.**
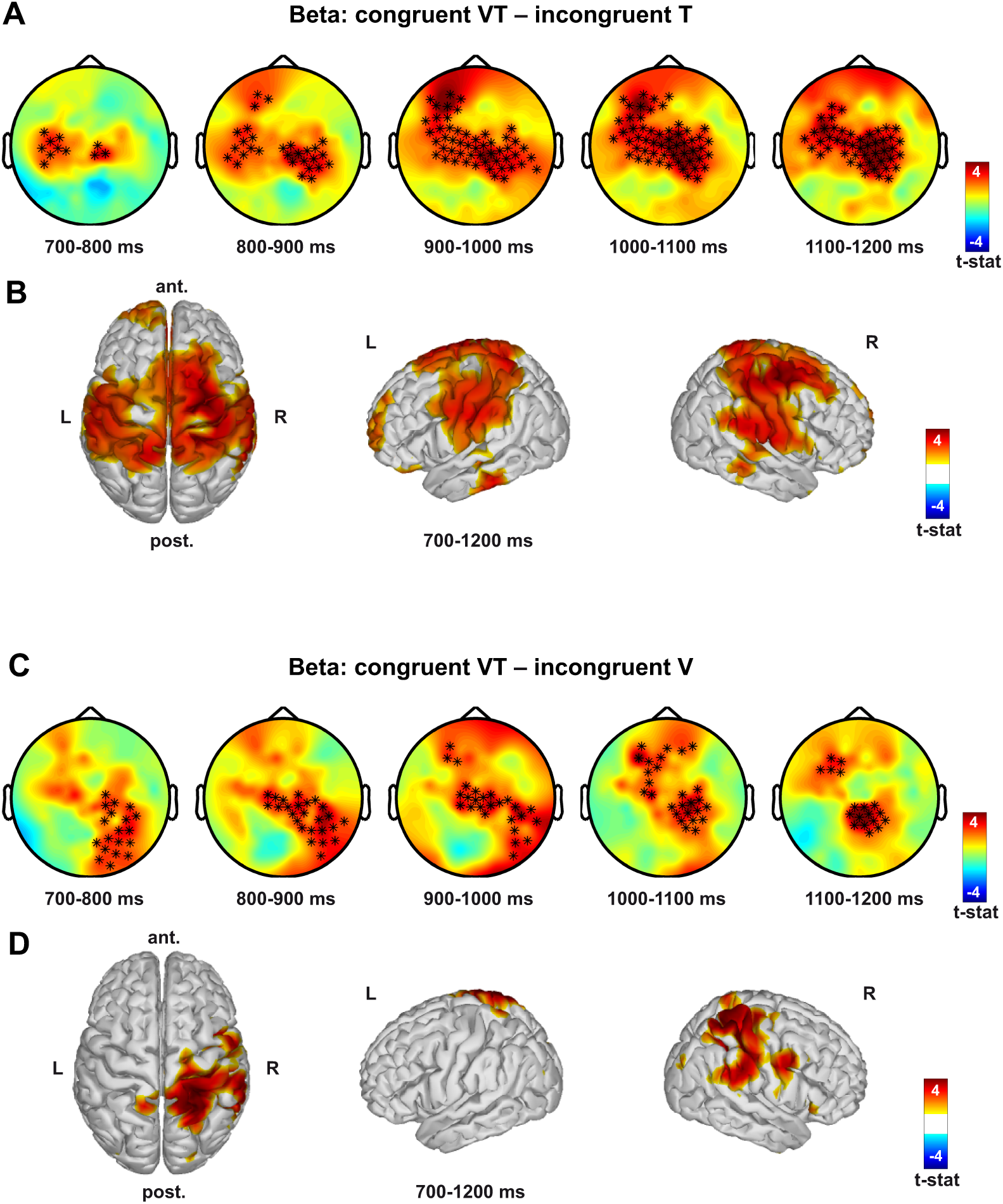
Comparison of beta-band responses in the *detection task*. **(A)** Topographies of the difference (shown as t-values) in beta-band power (13-30 Hz) between congruent VT and incongruent T target trials in 100 ms bins between 700 and 1200 ms after stimulus onset. A cluster (*p* = 0.006) in central scalp locations is apparent. (**B**) Statistical differences (shown as t-values) for the source level comparison of beta-band responses between 700 and 1200 ms for congruent VT and incongruent T target trials. T-values are masked using FDR correction (alpha = 0.1) to account for multiple comparisons (across voxels). Maxima are located in right-hemispheric premotor cortex, SMA and supramarginal gyrus, as well as left prefrontal regions. (**C**-**D**) Comparison of beta-band responses on congruent VT and incongruent V target trials. Presentation of results is analogous to (A) and (B). The cluster in (C) is significant with *p = 0.007*. Maxima of source level contrasts in (D) were found in right-hemispheric somatosensory association cortex and supramarginal gyrus. Asterisks in the topographies mark the channels belonging to the cluster showing significant difference between conditions.

Looking at the topography of late beta-band activity averaged over conditions (Figure 2C) it becomes evident that the observed differences related to crossmodal congruence are due to differences in right-hemispheric power decreases between conditions as well as beta-band rebound phenomena that are restricted to left-hemispheric regions.

#### 3.2.2 Congruence evaluation task

For the *congruence evaluation task*, power values were also compared across the whole post-stimulus period within the theta-, alpha- and beta-frequency ranges to assess differences related to visuotactile congruence.

Whereas no differences between congruent and incongruent stimulus pairs were observed for the alpha- and beta-band, a significant cluster in the theta-band was apparent (Figure 4A, *p* = 0.007) with higher power values being associated with incongruent pairs. This cluster shows a fronto-central distribution and evolves between 400 and 1100 ms after stimulus onset. Source statistics for the comparison of congruent and incongruent stimulus pairs in this time interval showed difference maxima to be located in right-hemispheric premotor cortex, prefrontal cortex, the ventral part of the cingulate cortex, and left superior temporal gyrus (Figure 4B).

**Figure 4.**
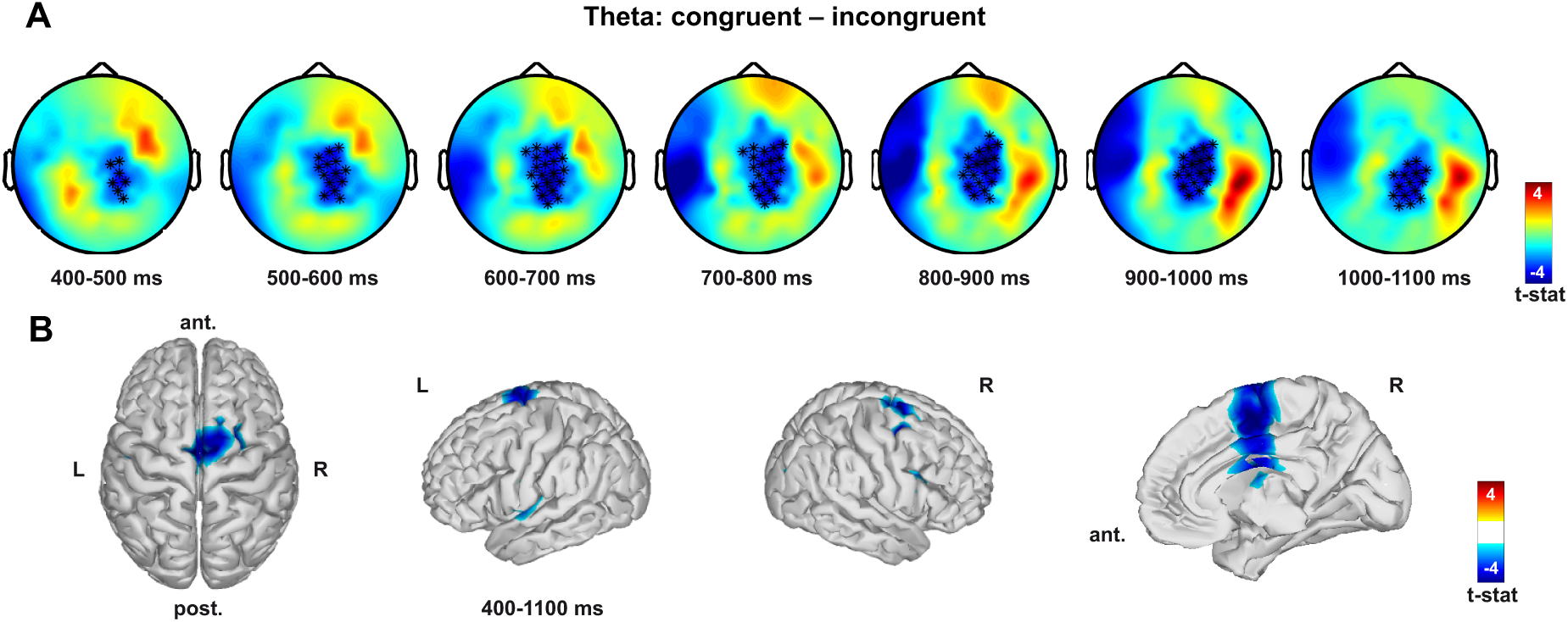
Comparison of theta-band responses in the *congruence evaluation task*. (**A**) Topographies of the difference (shown as t-values) in theta-band power (4-7 Hz) between congruent and incongruent trials in 100 ms bins between 400 and 1100 ms after stimulus onset. A cluster (*p* = 0.007) in fronto-central scalp regions is apparent. (**B**) Statistical differences (shown as t-values) for the source level comparison of theta-band responses on congruent and incongruent trials between 400 and 1100 ms. T-values are masked using FDR correction (alpha = 0.1) to account for multiple comparisons (across voxels). Maxima are located in right-hemispheric premotor cortex, prefrontal cortex, the ventral part of the cingulate cortex, and left superior temporal gyrus. Asterisks in the topographies mark the channels belonging to the cluster showing significant difference between conditions.

#### 3.2.3 Comparison between the detection and congruence evaluation tasks

To investigate global between-task differences in neuronal activity associated with distinct cognitive demands, we compared theta-, alpha- and beta-band specific responses (raw power values) for the *detection* and the *congruence evaluation task* in an interval directly preceding stimulus presentation (− 300 ms to 0, in steps of 100 ms). Data were averaged across conditions within the two tasks (trial numbers were matched for every participant) and thereafter compared between the tasks. Significant differences in the pre-stimulus activity were found in the alpha- and beta-frequency ranges. Employing a cluster level randomization approach, we found pre-stimulus alpha-band power to be reduced for the *detection task* as compared to the *congruence evaluation* task, resulting in a broadly distributed negative cluster (*p* = 0.003, Figure 5A).

**Figure 5.**
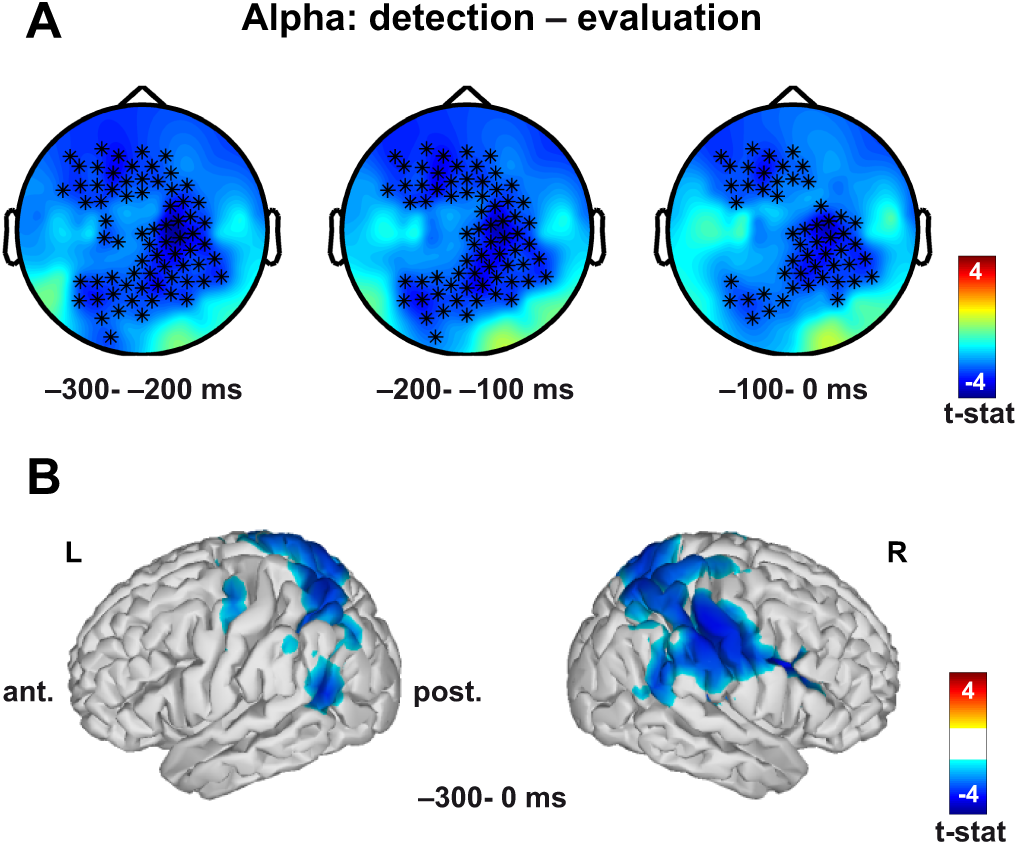
Comparison of alpha-band pre-stimulus activity between the *detection* and the *congruence evaluation task*. (**A**) Topographies of the differences (shown as t-values) in alpha-band power (8-13 Hz) in a time interval from of − 300 ms to 0 (onset of the visual and tactile stimuli) in steps of 100 ms. A broadly distributed, significantly negative cluster (*p = 0.003*) is apparent. (**B**) Maxima of source level differences for the comparison of anticipatory alpha-band power between the two tasks. T-values are displayed using a t-threshold of ± 2.1314 (corresponding to the 2.5^th^ and 97.5^th^ percentiles of the t distribution with 15 degrees of freedom). Maxima of source level differences are located in right-hemispheric premotor cortex and SMA, as well as the supramarginal gyrus. Asterisks in the topographies mark the channels belonging to the cluster showing significant difference between tasks.

Differences in anticipatory alpha-band activity were mainly located in right-hemispheric premotor cortex and SMA, as well as the supramarginal gyrus (Figure 5B).

Contrasting pre-stimulus beta-band responses (13 to 30 Hz) between *detection* and *congruence evaluation* also resulted in a negative cluster (*p* = 0.004), the spatial characteristic being more focal and distributed in right-hemispheric regions (Figure 6A). Also in the beta-frequency range, pre-stimulus power was reduced for the *detection task*.

**Figure 6.**
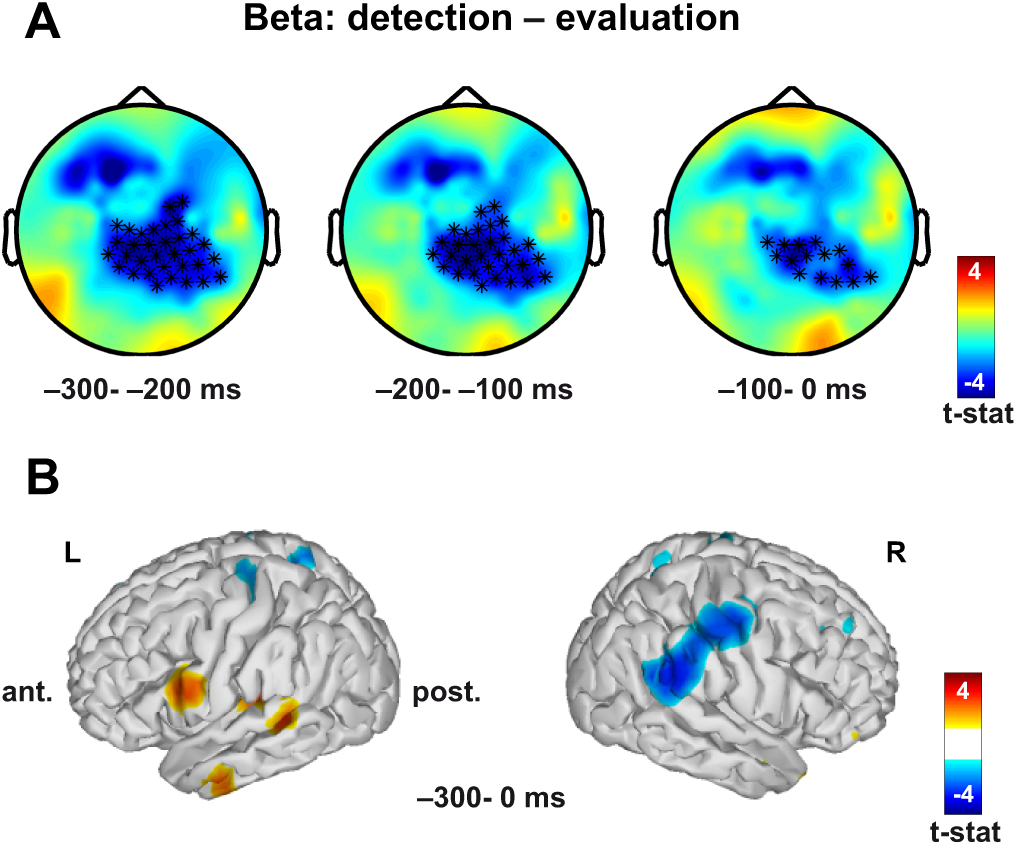
Comparison of beta-band pre-stimulus activity between the *detection* and the *congruence evaluation task*. (**A**) Topography of the differences (shown as t-values) in beta-band power (13-30 Hz) in a time interval from − 300 ms to 0 (onset of the visual and tactile stimuli). A significantly negative cluster (*p* = 0.004) is apparent in right-hemispheric scalp regions. (**B**) Maxima of source level differences for the comparison of anticipatory beta-band power between the two tasks. T-values are displayed using a t-threshold of ± 2.1314 (corresponding to the 2.5^th^ and 97.5^th^ percentiles of the t distribution with 15 degrees of freedom). Maxima of source level differences are located in the right supramarginal gyrus and somatosensory association cortex. Asterisks in the topographies mark the channels belonging to the cluster showing significant difference between tasks.

Contrasts for pre-stimulus beta-band activity between the two tasks corresponding to the negative cluster in sensor space peaked in the right supramarginal gyrus and somatosensory association cortex (Figure 6B). Positive differences were found in the left middle temporal gyrus and left premotor cortex (Figure 6B).

## 4. Discussion

In the present study we investigated behavioral and oscillatory signatures of visuotactile stimulus congruence effects by means of a crossmodal pattern matching paradigm involving two different tasks. On a behavioral level, we found evidence for stimulus-congruence related enhancement in performance, replicating our previous findings (Göschl et al., 2014).

Within the *detection task*, we found differences in oscillatory dynamics associated with pattern congruence in the beta-band between 700 and 1200 ms after stimulation, with congruent visuotactile stimulation being associated with higher power values. In the *congruence evaluation task* in contrast, we observed incongruent pattern combinations leading to augmented power in the theta-band in an interval from 400 to 1100 ms.

Comparing baseline activity (− 300 ms to 0) between the *detection* and the *congruence evaluation tasks*, differences in neuronal activity were apparent in the alpha- and beta-frequency ranges showing power values to be less pronounced for the *detection task*. In the following, we discuss stimulus congruence-related as well as task-related effects in our visuotactile matching paradigm in detail.

### 4.1 Detection task

Using stimulus material comparable to our study, the relevance of oscillatory brain activity in mediating multisensory interactions has been shown before (Bauer et al., 2009; Bauer et al., 2012; Kanayama and Ohira, 2009; Kanayama et al., 2012). Here, we add to the existing literature by showing that the crossmodal relation of stimuli presented in two sensory modalities is critical for performance and that congruence-related behavioral gains are related to low frequency oscillatory activity in the beta-band.

Beta-band activity in a time interval directly preceding participants’ response differed significantly between congruent and incongruent visuotactile stimulus pairs. There is evidence that beta-band activity is related to multisensory processing (e.g., Senkowski et al., 2006; Schepers et al., 2013) but its role in mediating crossmodal congruence effects is unclear. Differences in late beta-band power found in the current experiment show maxima in premotor cortex, SMA, somatosensory association cortex, supramarginal gyrus and prefrontal regions. Therefore, we hypothesize that processes of perceptual decision making may be reflected in these activation differences. Recent work by Donner and colleagues (2007, 2009) has linked beta-band activity to choice behavior in a visual motion detection task and observed that performance-predictive activity is expressed in posterior parietal and prefrontal cortices. Given the match of differences in behavioral performance for congruent VT targets, incongruent V and incongruent T targets and late beta-band power for the different conditions, we suggest that beta-band power may be linked to decision making also in the current study. In this sense, congruent visuotactile stimulus pairs as compared to the incongruent target cases might be viewed as stronger sensory evidence for an upcoming decision and the corresponding motor response (pressing the ‘target’ button). This choice-related activity could be reflected in beta-band power (Donner et al., 2007; Donner et al., 2009).

Beta-band activity in central regions might also relate to processes of response inhibition. Thus, increased power for congruent trials – that were associated with better performance – could reflect a higher demand for inhibiting the stronger urge to respond. Differences in task difficulty within the *detection task* complicate the interpretation of spectral differences. Visual targets were detected more easily than tactile ones but still, congruent stimulation (visual and tactile targets appearing together) was associated with the best performance. This result led us to conclude that there is behavioral facilitation related to crossmodal stimulus congruence going beyond differences in target detection between the two modalities. Differences in oscillatory signatures only partly reflect this relationship. While we do find stimulus-congruence related differences in beta-band power between congruent VT and incongruent T targets, as well as between congruent VT and incongruent V targets, we do not observe differences between incongruent T and incongruent V targets. Therefore we conclude that the observed differences in the beta-band not solely mimic task difficulty, but rather relate to visuotactile stimulus congruence facilitating crossmodal integration and detection performance.

The prominent involvement of the supramarginal gyrus is in line with work on tactile texture and pattern discrimination (Hegner et al., 2010) and consistent with a recent study assigning this region a key role in visuotactile integration (Quinn et al., 2014). Enhanced power responses located in the supramarginal gyrus for congruent crossmodal stimuli could point to inter-sensory facilitation effects in pattern discrimination resulting from visuotactile congruence and lead to improved detection performance.

### 4.2 Congruence evaluation task

Within the *congruence evaluation task*, differences in oscillatory dynamics between matching and non-matching visuotactile pairs were confined to the theta-band. In general, the modulation of theta-frequencies in the integration of features across sensory modalities is in agreement with previous reports (van Ackeren et al., 2014). For the comparison of congruent and incongruent stimulation, theta-band power has been shown to be more pronounced for the incongruent case which in turn has been linked to processes of conflict monitoring and conflict resolution, respectively (Cohen and Donner, 2013; Cohen and Ridderinkhof, 2013; Kanayama and Ohira, 2009). Similarly, we find theta-band power to be stronger for stimulus pairs that are incongruent across sensory modalities – thereby extending the definition of ‘congruence’, which mostly refers to spatial proximity. The observed effect is located in premotor cortex and prefrontal and cingulate cortices, comparable to previous reports on conflict processing (e.g., Cohen and Ridderinkhof, 2013). Of note, we only observe differences in theta-band activity for the *congruence evaluation task*, where the crossmodal relation of the two stimuli was explicitly task relevant. If higher theta-band power for incongruent trials relates to higher conflict, one might expect this relation to be also mirrored in the behavioral data. In fact, we only observe a trend (*p* = 0.06) for incongruent stimulus pairs being associated with lower performance, demanding a cautious interpretation of the behavioral relevance of the observed power effects. Nonetheless, data from a previous study with 39 participants (Göschl et al., 2014) show significant behavioral benefits for congruent stimulation in the *congruence evaluation task*.

### 4.3 Comparison between the detection and congruence evaluation tasks

For the comparison of neuronal activity between the *detection* and the *congruence evaluation task* we focused on the pre-stimulus period and found significant effects in the alpha- and beta-bands.

As suggested by the concept of functional inhibition by alpha-oscillations (Jensen and Mazaheri, 2010; Jensen et al., 2014; see also Klimesch et al., 2007), we hypothesized that differences in cognitive demands imposed by the two tasks would modulate preparatory oscillatory activity differentially in alpha-/beta-frequencies (see also Mazaheri et al., 2014). Contrasting alpha-band activity before stimulus onset for *detection* versus *congruence evaluation* indeed yielded significant differences showing a power decrease with broad, mainly right-hemispheric distribution. At the source level, maxima of these differences were located in right-hemispheric premotor cortex and SMA, as well as supramarginal gyrus. Interpreted within the framework of “gating by inhibition” (Jensen and Mazaheri, 2010; Jensen et al., 2014) decreased activity in anticipation of crossmodal stimulation could point to a higher engagement of these regions for the *detection task*. Similarly, differences in beta-band power before stimulus onset between the two tasks were mapped to a negative cluster in right-hemispheric regions, the spatial distribution being somewhat more focused. Source difference maxima in the supramarginal gyrus and somatosensory association cortex again suggest a more pronounced involvement of these regions in anticipation of crossmodal stimulation in the *detection task*.

The higher engagement of the supramarginal gyrus – an area that has previously been associated with visuotactile integration (Quinn et al., 2014) – for *detection* rather than *congruence evaluation* seems surprising. Whereas stimulus congruence only implicitly plays a role in the former task, the latter demands to explicitly evaluate the crossmodal relation of the two stimuli. On the other hand, cortical areas showing differential activation in the baseline period between the tasks – namely premotor cortex, SMA, somatosensory association cortex and the supramarginal gyrus – are those being crucially involved in differences within the *detection task* related to crossmodal stimulus congruence. Future work needs to further determine the relation of task demands and bottom-up stimulus congruence and their reflection in neuronal oscillations.

In the current experiment, we realized lateralized stimulus presentation in the investigation of crossmodal interactions. The question of whether the observed cortical activations result from this lateralized stimulation or rather from cortical asymmetry goes beyond the scope of the current work and remains to be clarified in future studies. However, the pronounced involvement of right-hemispheric cortical regions for the spatial pattern matching task used here is well compatible with experimental evidence on a dominant role of the right hemisphere in spatial processing (see for example Hegner et al., 2010).

### 4.4 Conclusions

The current study adds to increasing evidence that neuronal oscillations are involved in multisensory interactions. Specifically, oscillatory activity in lower frequency ranges (below 30 Hz) seems to be relevant for long-range communication that is crucial for crossmodal processing. Here we studied visuotactile interactions as a model for integration in distributed brain networks and evaluated crossmodal congruence in two different tasks using physically identical stimulation. We found different spectral signatures of congruence-related effects depending on distinct task demands. Cortical areas mediating congruence-related effects in crossmodal *target detection* – most importantly the supramarginal gyrus – also showed more engagement in the baseline comparison between the tasks.

## 5. Acknowledgments

This research was supported by grants from the German Research Foundation (SFB 936/A3/B6) and the European Union (ERC-2010-AdG-269716) awarded to A.K.E. and P.K. The authors thank Julia Diestel for the assistance in data recording, Till Schneider for the helpful discussions on previous versions of the manuscript and Guido Nolte, Arne Ewald and Peng Wang for their methodological counseling.

